# Inter-species stimulus enhancement: Herring gulls (*Larus argentatus*) read human behaviour during foraging

**DOI:** 10.1101/2022.11.18.517022

**Authors:** Franziska Feist, Kiera Smith, Paul Graham

**Affiliations:** School of Life Sciences, University of Sussex, Brighton, East Sussex, BN1 9QG, United Kingdom

**Keywords:** social learning, stimulus enhancement, herring gulls, kleptoparasitism, inter-species learning

## Abstract

Herring gulls are one of the few species that thrive in anthropogenic landscapes and their familiarity with people makes them an excellent target for studies of inter-species social cognition. Urban gulls pay attention to human behaviour in food-related contexts and we set out to investigate whether such cues can be used to redirect a gull’s attention to potential food items in their environment. Herring gulls were given free choice of two differently coloured anthropogenic food items in the presence of a demonstrator, who was either sitting still or pretending to eat food from an item that matched one of the secondary food items. We found that a demonstrator mimicking eating significantly increased the likelihood of an approach or peck. Furthermore, 95% of pecks were directed towards the secondary food item which colour-matched the demonstrator’s food item. The results show situation-dependent attentional modulation in gulls, whereby gulls are able to use human cues for stimulus enhancement and foraging decisions. Given the relatively recent history of urbanisation in herring gulls, this cross-species social information transfer is likely to be a by-product of the cognitive flexibility inherent in kleptoparasitic species.

## Introduction

The expansion of cities (Lowry, Lill and Wong, 2013) often negatively affects wildlife (Bateman and Fleming, 2012). However, some species thrive in anthropogenic landscapes and urban numbers can exceed population densities found in natural environments (Bateman and Fleming, 2012). This is true for the European herring gull (*Larus argentatus*) where, following the first instances of urban nesting in the 1940s, such colonies have spread across the UK (Rock, 2005), and despite overall population numbers declining (BirdLife International, 2018), urban populations continue to increase (Rock, 2005). As a result, many herring gulls are familiar with the presence of people and the foraging opportunities they provide.

Herring gulls are generalist predators that will feed on intertidal marine invertebrates and fish (Pierotti and Good, 1994), as well as human food items (Pierotti and Annett, 1991; Lato et al., 2021). This use of anthropogenic food sources indicates that urban gulls have adjusted their diet and foraging strategies to urban environments, an adaptation that may have been aided by their generalist foraging behaviours extending to kleptoparasitism, or food-stealing. Many gulls steal from both hetero- and conspecific individuals (Thompson, 1986; Busniuk et al., 2020), with occurrence depending on species, age, profit and the environment (Bertellotti and Yorio, 2000; Spencer et al., 2017; Goumas et al., 2019; Busniuk, Storey and Wilson, 2020). Such behavioural flexibility in foraging is suggested to be one of the factors contributing to high survival rates in novel environments (Spencer et al., 2017) but is likely to require specific cognitive adaptations.

Behavioural and cognitive flexibility have been shown to facilitate adaptation to human landscapes in other species (Bateman and Fleming, 2012; Plumer, Davison and Saarma, 2014; Lee and Thornton, 2021). Specifically, an increased capacity to acquire, store and process information about one’s environment is hypothesised to improve survival rate (Lee and Thornton, 2021). For food-stealing birds, success is said to depend on the ability to integrate and use information about the environment and other individuals, as this facilitates the selection of targets, approach strategy and the prediction of host-response (Morand-Ferron, Sol and Lefebvre, 2007). Furthermore, kleptoparasites generally have larger brain sizes than their hosts, an association that remains significant even after controlling for variations in juvenile development period (Morand-Ferron, Sol and Lefebvre, 2007).

Herring gulls, specifically, show evidence of social learning via the acquisition of information from observation of hetero-/conspecifics (Damas-Moreira et al., 2018), and their long life span and four-year juvenile period (O’Hanlon and Nager, 2017) provide ample opportunities for extensive social learning. In social learning and foraging, gulls rely extensively on conspecifics, and often obtain food after watching others flock to a food source (Goumas, Boogert and Kelley, 2020a). This suggests that gulls make use of social learning to learn the relationship between a stimulus and a demonstrator, which may then allow them to respond similarly in the same scenario (Hoppitt and Laland, 2008). Interestingly, in urban populations there is evidence that herring gulls have also learned to obtain foraging information from humans. Recent studies have found gulls are able to adapt their foraging behaviour to human activity patterns (Spelt et al., 2021), increase their attention towards a person in possession of food (Feist and Graham, 2022), and pay attention to behavioural cues such as gaze (Goumas et al., 2019; Goumas et al., 2020b) and to which items have been handled (Kelley, Boogert and Goumas, 2020a). This shows that gull populations have the cognitive and behavioural flexibility to adapt foraging behaviour to human cues.

We further study the cognitive abilities of gulls by asking if they can not only pay attention to humans and human food items, but also to transfer knowledge from human behaviour onto secondary food items via social observation. Using a stimulus enhancement assay, we ask if the specific details, in our case colour, of a food item being consumed by a human can influence the choice of secondary food items by herring gulls, so called stimulus enhancement.

## Methods and Materials

### Study site

Data was collected during daylight hours (7:00-16:00) along the Brighton beachfront, UK (50.8193° N, 0.1364° W), from May-June 2021 and March-May 2022. Differing weather conditions only affected gull presence but not behaviour (Feist and Graham, 2022), however low tide periods were avoided as gulls engage more in natural foraging and were consequently less likely to engage with the experiment. Furthermore, data was only collected on weekdays to minimise pedestrian disturbance.

### Data collection

Single individual herring gulls and groups of less than five were approached after assessing the area to ensure there were no dogs or pedestrians moving in the immediate vicinity. Two Walkers brand crisp packets, one blue, one green, taped to (15 cm × 20 cm) ceramic tiles were laid out, spaced approximately 1.5 m apart and roughly 5-10 m away from the gulls. The left-right position of the packets was alternated to control for any side bias. The experimenter retreated and sat on the ground approximately 5 m from the crisp packets. Subjects were recorded on an iPhone XS or Honor 20 mounted to a LINKCOOL tripod (Fig. 1A). Recording was stopped when a gull pecked at one of the food items or after five minutes had passed, unless a gull was approaching at the five minute point. No individuals were forced to engage with the experimenter, and all gulls had the freedom to remove themselves at any time during trials by walking or flying away. The experimenter adhered to guidelines set by the Association for the Study of Animal Behaviour (ASAB) surrounding the treatment of animals in behavioural research and testing (Buchanan et al., 2012) and all necessary ethical approval was obtained from the University of Sussex.

**Figure 1.**
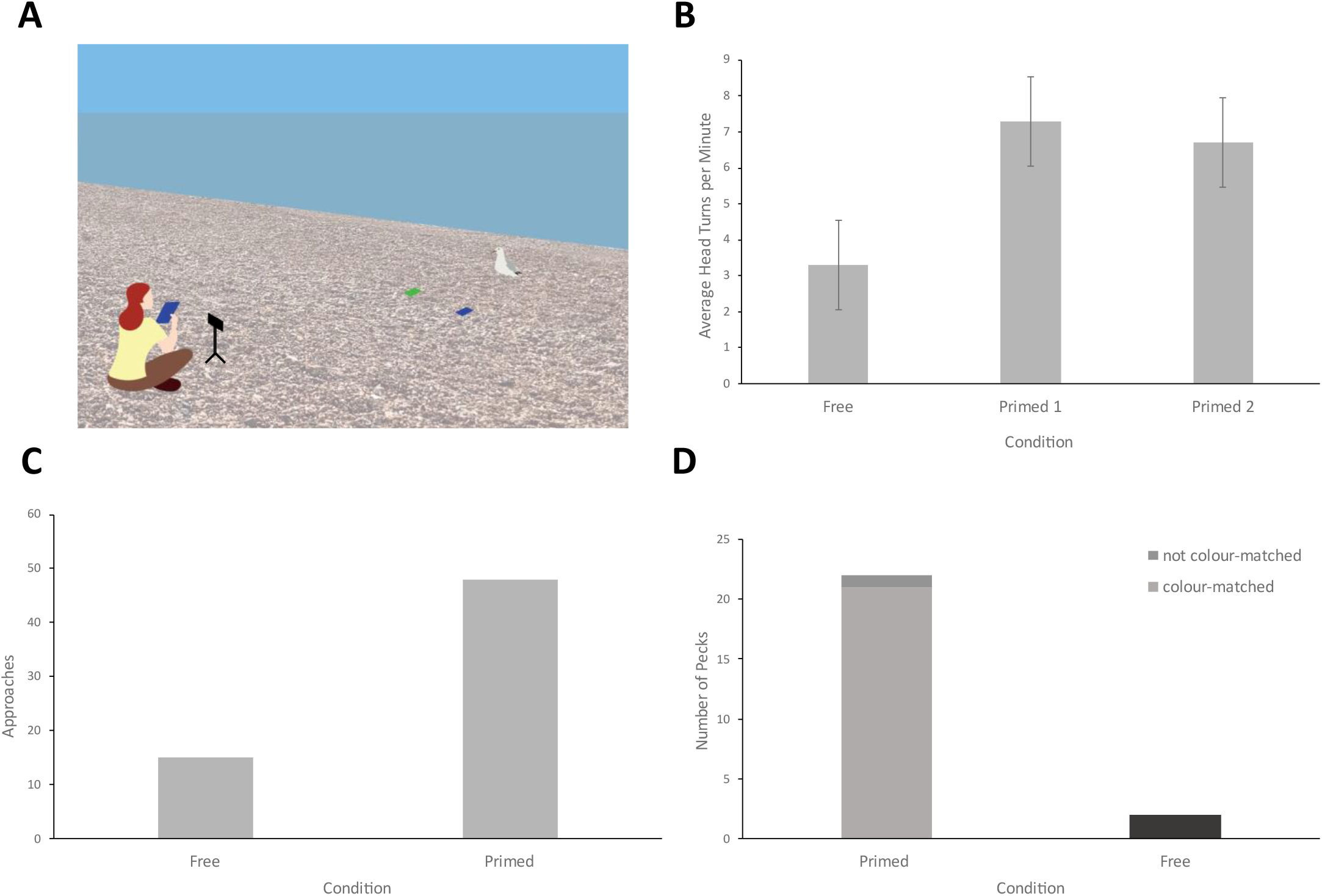
Gull attention and foraging behaviour is biased by human behaviour. **(A) Illustration of the experimental set-up**. After placing the crisp packets (blue and green square) in the vicinity of the target gull, the experimenter retreated behind the camera and either remained like that (FC) or retrieved their own packet of crisps (PC1 and PC2). **(B) Upregulation of attention during primed trials**. The average number of head turns per minute are significantly upregulated during PC (n=61) trials (X2=23.45, df=2, n=93, p<0.001) compared to FC (n=32) trials, with significantly more head turns in both the PC1 (W=840.5, p<0.001) and PC2 (W=412, p<0.001) condition. However, the difference between PC1 (n=38) and PC2 (n=23) was not significantly different (W=259, p=0.913). **(C) Approach instances are higher during primed trials**. Approaches happened significantly more often in PC (n=103) trials (X2=132.75, df=1, n=183, p<0.001), with 51.46% of birds approaching during primed choice compared to only 18.75% during free choice trials (n=80). **(D) Most pecks occurred during primed trials and colour-matched the hand-held object**. Pecks happened significantly more often in primed choice (n=103) compared to free choice (n=80) trials (X2=197.18, df=4, n=183, p<0.001), with a total of 21.36% and 2.5% of birds pecking the crisps bags on PC and FC trials respectively. Additionally, pecks were strongly biased to the packet which colour-matched the primary food item held by the human experimenter in PC trials (95.45% of pecks; W=0, n=22, p<0.001).

We measured the gull’s behavioural responses to the experimenter and secondary stimuli in three different conditions. The first condition, free condition (FC), involved the experimenter retreating and sitting neutrally, holding a fixed gaze to control for human gaze cues (Goumas et al., 2019). The second condition, primed condition 1 (PC1), involved the experimenter retrieving a blue crisp packet from their bag and eating from it, whilst the third condition, primed condition 2 (PC2), involved the experimenter retrieving and eating from a green crisp packet.

### Video Analysis

Recordings were uploaded into the Behavioural Observation Research Interactive Software (BORIS) program (Friard and Gamba, 2016) and an ethogram of head turns and approaches was created from video recordings, while pecks were simply recorded as “yes” or “no”. A head turn was recorded if the gull turned its head towards the experimenter, and the average number of head turns per minute per individual was calculated. In order for an individual to be included in the analysis it had to be present in the video for 30 seconds or more, or peck at one of the provided crip packets. Approaches were defined as the gull moving from its original position to at least halfway towards the food items, and a peck was recorded if the gull’s beak came into contact with a crisp packet.

The age of the gulls was recorded as juvenile or adult based on their plumage. Any individual with brown markings, regardless of any grey wing feathers observed, was recorded as a juvenile. Using the data from 2022 only, the frequency of each attention marker was compared between adults and juveniles. The directness of approaches was ranked from 1-5 by an independent observer, with 1 being least direct and 5 being most direct.

### Statistics

Data was logged in Microsoft Excel and loaded into R studio version 4.0.3 (R Core Team, 2022). Normality was assessed with a Shapiro Wilk test, and a Kruskal Wallis test was used to look for differences between the average number of head turns per minute in the food conditions (FC, PC1 and PC2). A post hoc, pairwise Mann Whitney U test was performed to assess the impact of each condition individually, and to look at the impact of age on the average number of head turns. Furthermore, we used a Chi-squared test to assess the impact of age and condition on the proportion of birds that approached or pecked at one of the crisp bags. Lastly, Mann Whitney U tests (Thomas, 2015) were used to assess the impact of food condition on the directness of approaches.

## Results

### Human behaviour impacts gull attention

We first aimed to establish the impact of human food cues on three markers of attention: head turns, approaches and pecks. All three were increased during PC1 and PC2 compared to FC (Fig. 1B-D). The average number of head turns per minute (Fig. 1B) differed significantly between conditions (X2=23.45, df=2, n=93, p<0.001), with significantly more head turns in both the PC1 (W=840.5, p<0.001) and PC2 (W=412, p<0.001) when compared to FC. The difference between PC1 and PC2 was not statistically significant (W=259, p=0.913), which confirms that the colour of crisp packet did not affect attention, thus we subsequently combined data from the two primed conditions (PC).

Approaches (Fig. 1C) occurred significantly more often in PC trials (X2=132.75, df=1, n=183, p<0.001), with 51.46% of birds approaching during primed choice compared to only 18.75% during free choice trials. Similarly, pecks (Fig. 1D) happened significantly more often in primed choice compared to free choice trials (X2=197.18, df=4, n=183, p<0.001), with a total of 21.36% and 2.5% of birds pecking the crisps bags on PC and FC trials respectively.

We further analysed markers of attention by age. The average number of head turns did not differ between age categories, irrespective of condition. In FC trials, we recorded an average of 3.61 head turns per minute for adults and 3.09 for juveniles (W=762, p=0.2). Similarly, in the primed condition the mean was 7.47 and 6.13, respectively (W=352.5, p=0.09). In contrast, the proportion of approaches (X2=108.32, df=4, n=93, p<0.001) and pecks (X2=114.1, df=4, n=93, p<0.001) differed significantly with age during PC trials, with 74.19% of approaches and 94.74% of pecks coming from adults, which made up 59% of the target population.

### Human food cues affect gull attention

We further asked if gulls observe the details of human behaviour and use that information to redirect their attention towards a specific, secondary stimulus. Gulls approached a secondary food item (crisp packet) more directly in primed choice trials compared to free choice trials (W=93, n=59, p<0.001) and approaches that resulted in a peck (50%) were strongly biased to the packet which matched the food item held by the experimenter (95.45% of pecks; W=0, n=22, p<0.001).

Whilst our results clearly indicate that gulls will frequently show interest in an anthropogenic food item on the ground, the same objects are often ignored. During free choice trials (n=80), a total of 15 gulls approached (18.75%), but only two (2.5% of total or 13.33% of approaching birds) pecked at the crisp bags on the ground. In comparison, during primed choice trials (n=103), 53 gulls approached (51.46%) and ten (9.71% of total or 18.87% of approaching birds) ignored the crisp bags on the floor and walked past them, approaching the experimenter instead.

## Discussion

Our primary research question was whether gulls could pay attention to human food cues and use those to influence choices towards secondary objects. As expected, we confirmed that gull attention is captured when humans are seen eating. However, we also observed that when gulls approach and peck at a food stimulus, they choose the item that colour-matches that held by the experimenter. As expected (Goumas et al., 2019), birds that approached represented a subset of gulls which are probably specialised human-food stealers. Additionally, whilst some birds pecked at the secondary food stimulus, others approached the experimenter directly, reflecting a balance of stimulus enhancement and local enhancement. We discuss these results in terms of gull cognition and learning.

Whilst comparative attempts to categorise (or rank) the cognition of gulls relative to other animals have been inconclusive (Beck, 1982; Benjamini, 1983; Obozova, Smirnova and Zorina, 2012; Zorina and Obozova, 2012), it is clear from naturalistic foraging tasks that gulls have rich and flexible behavioural repertoires, suggestive of a high level of cognition. For instance: Observations of herring gulls dropping shells demonstrate persistence, concentration and mental representations of the distribution of drop sites (Beck, 1982); Systematic food handling, such as removing large pincher claws before returning to water to dip prey prior to consumption reflects skill and purposefulness (Beck, 1982). Furthermore, urban herring gulls are capable of object tracing (Kelley, Boogert and Goumas, 2020a) and Glaucous-winged gulls (*Larus glaucescens*) have been shown to be quick social learners (Obozova, Smirnova and Zorina, 2011). We further extend the understanding of gulls’ cognitive capabilities, in terms of objects and social cues. Our subjects show the ability to read human behaviour and make a connection between a stimulus on the ground and that held by the experimenter. Inference that the hand-held bag is identical to one of the bags on the ground allows gulls to make foraging choices influenced by their understanding of human behavioural cues.

An increased attention towards human cues in a food-related context is frequently seen in domesticated animals. Both dogs (*Canis lupus familiaris*) and horses (*Equus caballus*) can find hidden food by paying attention to human cues (Call et al., 2003; McKinley and Sambrook, 2000; Proops and McComb, 2010; Proops, Walton and McComb, 2010; Krueger et al., 2011). Similarly, while human gaze cues are not sufficient for goats (*Capra hircus*), human touching and pointing cues can be used to locate food items (Kaminski et al., 2005). Urban gulls may not be domesticated, but their history of adaptation to anthropogenic environments and frequent interactions with humans may have resulted in an increase of attention to, and understanding of, human food cues.

An alternative explanation is that kleptoparasitism, rather than urbanisation, is the driver for cross-species reading of behaviour. Wild brown skuas (*Catharacta antarctica* ssp. *lonnbergi*), another kleptoparasitic seabird, prefer food that has previously been handled by an experimenter (Danel et al., 2022) which raises questions as to whether frequent contact with humans is a prerequisite for the exploitation of human cues, or simply a facilitator, as some animals may possess a general tendency for paying attention to heterospecific cues.

Whether or not our observed gull behaviour is due to increased contact with anthropogenic scenarios, the pattern of results with age strongly suggests that learning plays an important role. Gulls learn to identify valuable food sources from observing conspecifics (Goumas, Boogert and Kelley, 2020a) and juvenile gulls learn and improve foraging skills as they mature (Gamble, 2000). Such stimulus enhancement, where observed object-focussed behaviour increases the salience of an object, may explain most of our results. Juveniles and adults may pay an equal amount of attention to human food cues, evidenced by head turns, but adults are more likely to take this interest further by approaching or pecking at the food item. Juveniles may lack the skill or boldness to launch a kleptoparasitic attack on a person, a notion that is supported by Källander (2000) who found that juvenile kleptoparasitic success in black headed gulls (*Larus ridibundus*) increased over time. Similarly, Monaghan (1980) found that the number of adult herring gulls foraging at tip sites was significantly higher than the proportion of adults in the population, suggesting that interest in human food items may be higher in adults. If the development of foraging skill depends on social learning, this would be another marker of cognition in gulls (Morand-Ferron, Sol and Lefebvre, 2007).

Interestingly, some individuals were observed moving to the location of the experimenter rather than the secondary food items, suggesting those gulls are motivated by local enhancement (Hoppitt and Laland, 2008). In our case, this would mean that gulls approached because the experimenter eating at that location increased the attractiveness of a point in space, or of the experimenter themselves. This process has been reported in yellow legged gulls (*Larus michahellis*) and Cory’s shearwater (*Calonectris borealis*) when identifying suitable foraging areas (Sol, Arcos and Senar, 1994, Bastos et al., 2020), and its presence in gulls has been suggested by Goumas, Boogert and Kelley (2020a).

The distinction between gulls acting under stimulus enhancement versus local enhancement may be part of a broader heterogeneity in the anthropocentric behaviours in urban herring gull populations. Our data agrees with existing literature which states that only around a quarter of gulls will launch kleptoparasitic attacks on humans (Goumas et al., 2019). This is consistent with human-gull interactions requiring skill and boldness and there being a subset of human-foraging specialists within the population.

## Data availability

The data are available upon request.

