## Supplemental Figure 1 for "Inter-species stimulus enhancement: Herring gulls (*Larus argentatus*) read human behaviour during foraging"

### Supplemental Material

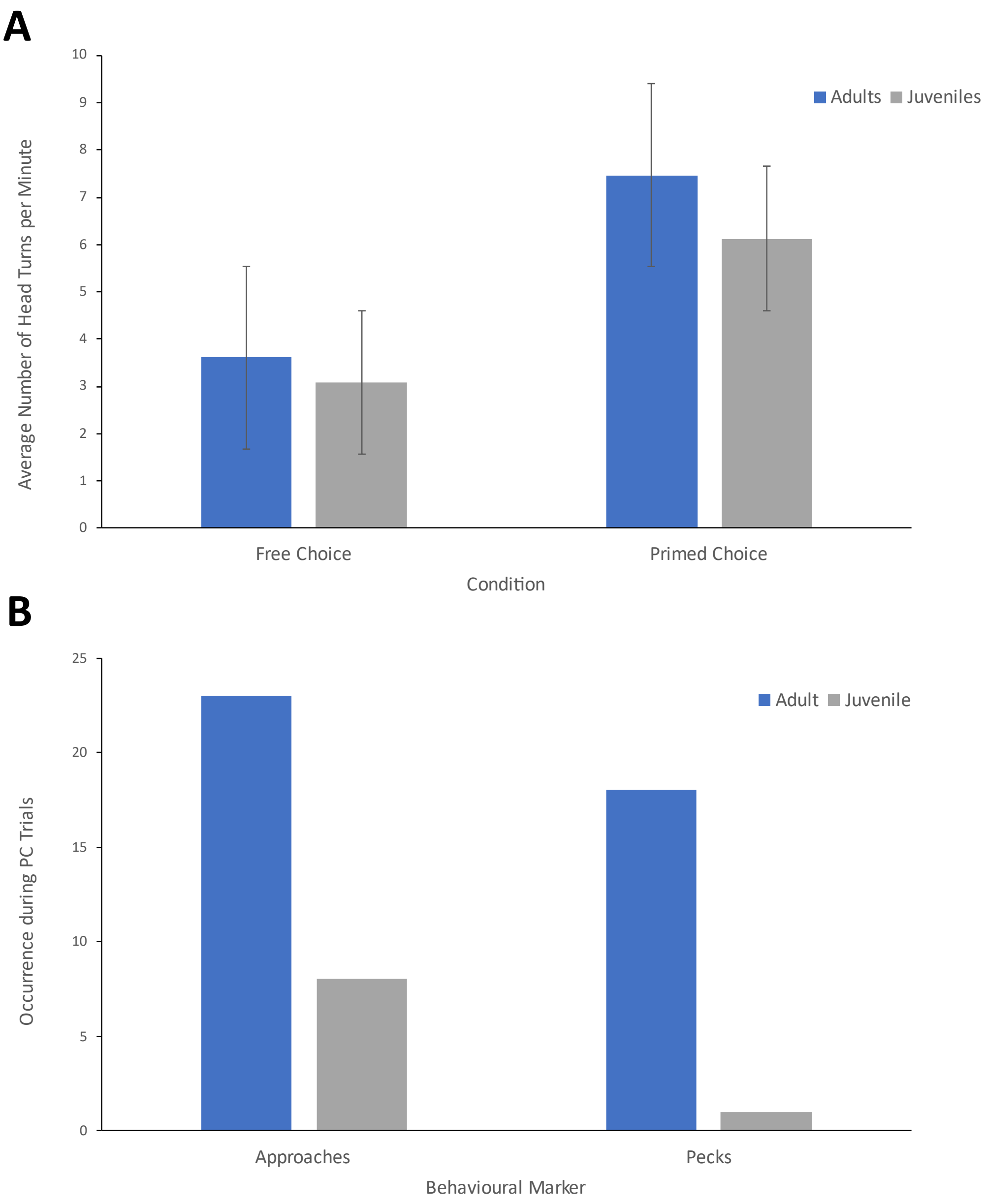


**Supplemental material S1. Age differences in occurrence of attentional markers. (A)** The average number of head turns per minute did not differ significantly between age groups, irrespective of trial condition. In FC trials (n=32), we recorded an average of 3.61 head turns per minute for adults and 3.09 for juveniles (W=762, p=0.2). Similarly, in the primed condition (n=61) the average was recorded to be 7.47 and 6.13, respectively (W=352.5, p=0.09). **(B)** The number of approaches (X2=108.32, df=4, n=93, p<0.001) and pecks (X2=114.1, df=4, n=93, p<0.001)

differed significantly with age during PC (n=61) trials, with 74.19% of approaches and 94.74% of pecks coming from adults, which made up 59.02% of the target population.
